# Leadership in PhD (LeaP): a longitudinal leadership skill building program for underrepresented biomedical research trainees

**DOI:** 10.1101/2022.09.11.507461

**Authors:** Mali D Doles, Ji Yun Kang, Linda M Scholl, Jason D Doles

## Abstract

Increasing diversity in the biomedical research workforce is a critical national issue. Particularly concerning is the lack of representation at more advanced career stages/in leadership positions. While there are numerous institutional initiatives promoting professional research skills (i.e. grant writing, presenting, networking) for underrepresented (UR) PhD trainees, there are comparatively fewer opportunities for leadership development. We present a blueprint for Leadership in PhD (LeaP), a cohort-based program aiming to equip UR biomedical research trainees with skills to succeed as academic, industry, and community leaders. In contrast to intensive short-term programs or workshops, LeaP is a longitudinal 4-year experience with an emphasis on self-directed and experiential learning. First year trainees receive foundational didactic instruction on core leadership concepts coupled with facilitated peer discussions and one-on-one coaching support. We outline a program evaluation framework that assesses student learning, satisfaction, and program efficacy. Evaluation data from the inaugural year is presented and discussed.

## 1 Introduction

Despite efforts to diversify the composition of the scientific workforce, stark disparities remain (1). A recent analysis conducted by the Pew Research Center in 2021 found that Black and Hispanic workers remain underrepresented in the science, technology, engineering and math (STEM) workforce compared with their share of all jobs (2). This disparity is acutely evident in the life sciences where Black and Hispanic workers comprise 6% and 8%, respectively, of the workforce despite overall representation across all jobs being 11% and 17%. Representation of women in the life sciences has fared better – in 2019 women comprised 48% of the life sciences workforce compared to 34% in 1990. Intersectional inequalities, however, compound individual deficits in representation. These inequalities are evident when considering the number of minority women in science, and further, when examining performance-based metrics including sub-discipline funding rates and scholarly impact (3). Perdurance in academic science also lags in minoritized groups. A recent study by Lambert and colleagues surveying postdoctoral fellow career choices found that academia-bound UR postdocs have significantly lower confidence in securing funding, feelings of self-worth, and research career self-efficacy. Exit rates of minoritized post-docs from research/academia were also significantly higher than their white/majority counterparts (4).

Representational diversity and equity concerns persist for those fortunate to secure research intensive faculty-level positions (5, 6). Bias within the promotion and tenure process coupled with a lack of cultural sensitivity and social awareness regarding the professional and personal needs of UR faculty has resulted in substantial attrition. Women, for example, reached a remarkable milestone in 2003 when rates of medical school matriculation first hit the 50% mark. Despite that achievement, the proportions of women associate professors (37%) and professors (25%) in medicine in 2020 indicate that academia is not capitalizing on those gains in student interest in pursuing medical and science-based careers (7). Due to numerous individual and systemic roadblocks hindering progress through academic rank, the number of women and underrepresented faculty in leadership positions remains low. According to a 2017 overview of university presidents reported by the American Council on Education, women comprised 30% of all University president positions. Further, within the already slim proportion of women presidents, only 9% identify as Black women, and only 4% as Hispanic women – compared to the roughly 83% who identify as White (8). A closer look at these numbers further reveals another disturbing trend in that minority women who do attain senior leadership are often serving at lower-ranked, less prestigious schools with small endowments, and limited resources for research and advanced scholarship (8). These intersectional inequities are not limited to the University presidents. Across all research and health sciences jobs, women hold 52% of reported leadership positions, while racial/ethnic minorities represent 11% of leadership, and Black/Hispanic women only 3% (9).

Addressing representational inequities in biomedical and senior academic leadership promotes individual fairness and justice, and if done in conjunction with rigorous training and support, could tackle a much larger issue of culture and climate concerns in academia. Many from minoritized groups report difficulties penetrating, enduring, and thriving within the privileged (i.e. older, white, and male) academic ecosystem. Diversifying leadership would inherently promote and foster professional networking and development of underrepresented trainees/junior fellows/faculty who have shared life experiences and seek role models/inspiration. Indeed, representational diversity is known to be a strong catalyst for career progression within underrepresented communities, which further yields benefits for the overall enterprise (10). A further benefit of intentional leadership training is tackling the pervasive culture of ineffective leadership and poor mentoring in academia (11–14) – a state largely resulting from inadequate training, inconsistent continuous professional development, and little accountability. Racism, xenophobia, and microaggressions present additional barriers for UR trainees and contribute to heightened stress, demotivation and feelings of exclusion. LeaP supports students as they navigate these and other issues as a cohort, while honing critical resiliency skills.

Leadership development at academic research centers is not new. Many institutions, centers, and scientific societies have stepped up and begun addressing this critical societal need. Key programmatic activities include robust mentor training initiatives, resilience and wellness support for diverse trainees, networking opportunities, and targeted leadership development opportunities. Notable leadership from the National Institute for General Medical Sciences, the National Science Foundation and cross-institutional networks/alliances including the National Research Mentoring Network (NRMN) (15) and the Leadership Alliance (16) has seeded a movement of academics dedicated to improving culture/climate and fostering the development of diverse leaders (17). That said, significant gaps and opportunities remain. First, many (if not most) leadership initiatives target individuals at an advanced career stage – typical requirements include holding an advanced degree and/or an active or pending faculty appointment. Second, many programs employ an intensive, immersion-based approach – excellent for building knowledge and seeding skills but limited in the ability to support longitudinal skill perdurance and further development. Third, programs targeted to early-stage trainees, such as advanced degree candidates, tend to feature a defined curriculum that prioritizes knowledge building and group skills while providing fewer opportunities for individual skills exploration and development. With LeaP, we aimed to create a supportive and flexible learning environment for underrepresented PhD trainees to discover, explore, and refine leadership skills essential for leading an inclusive workplace. While LeaP does not have the ability to singlehandedly address systemic issues hindering UR advancement in academia, we contend that it does have significant value as it provides UR trainees with foundational skills to succeed and thrive as leaders in the biomedical sciences. Here, we outline a conceptual framework for LeaP, describe key programmatic features, present a summary and evaluation of year one, and discuss future directions for this work.

## 2 Program Overview

### I. Conceptual framework

The core mission of the LeaP program is to facilitate the development of knowledge, skills, and abilities essential for future biomedical research leaders in academia, industry, and society. The cohort-based model is intended to provide a safe and encouraging environment in which to engage in active self-discovery and peer-to-peer support. A guiding principle of the LeaP model is self-directed learning, a popular educational concept championed by many in the leadership development space, particularly in professional school and corporate/enterprise settings (18–21). In the context of LeaP, self-directed learning is promoted using a blended learning approach featuring didactic seminars, facilitated discussions, one-on-one coaching, and experiential opportunities. Since scholars engaged with LeaP are biomedical PhD trainees, concerted efforts were made to 1) provide scholars with experiences complementary to but distinct from thesis advisory committees or workshop-based professional development opportunities, 2) tailor contact hours to fit within the constraints of a rigorous PhD curriculum, and 3) support progressive leadership development throughout their biomedical PhD training.

### II. Cohort selection

One of our primary objectives was to foster development of leadership skills in trainees historically underrepresented in biomedical research. We utilized the National Institutes of Health (NIH) definition of underrepresented in the sciences and further widened the umbrella to include any student with a demonstrated commitment to diversity, equity, and inclusion in addition to leadership development. A modified version of the PhD application rubric was used – one that equally prioritized the following four areas: 1) leadership potential, 2) research potential, 3) industry, persistence and commitment to education/personal growth, and 4) academic performance/research experience. With applicant permission, LeaP candidacy was evaluated in parallel to the PhD admissions process. Eight students were selected for the inaugural LeaP cohort.

### III. Leadership competencies

The four-year LeaP experience centers around leadership competencies described in and assessed by the Occupational Personality Questionnaire (OPQ) developed by SHL Group Limited (22). While the OPQ is broadly used in the context of talent acquisition and management, Mayo Clinic has long partnered with SHL to leverage the OPQ for upper-level leadership preparation. Here, the OPQ identifies strengths, working styles, and opportunities to develop capability and unlock potential – thus setting the stage for individualized leadership coaching. We aimed to adapt and streamline this approach for LeaP participants. Ultimately, ten core leadership competencies were selected from the OPQ that served as the foundation for the didactic and discussion-based material covered in Year 1: 1) leading and deciding, 2) working with people, 3) adhering to principles/values, 4) presenting and communicating, 5) persuading and influencing, 6) creating and innovating, 7) planning and organizing, 8) adapting and responding to change, 9) coping with pressures and setbacks, and 10) relating and networking (22). Program activities were aligned to these competencies and are described in more detail below.

### IV. Programmatic components

The overarching structure of the LeaP curriculum involves iterative and progressive cycles of concept learning, facilitated peer discussion, self-reflection, and practice/implementation. This structure is reflected in the year-to-year plan; year one focuses on concept assimilation, cohort building, self-discovery, goal setting, and skill practice, year two on continued self-directed learning and skill implementation/practice (e.g. choosing study topics and materials, leading discussions, mentoring first year LeaP scholars, pursuing community projects, etc.), year three on individual or small group-based intramural (institutional) leadership experiences, and year four on tailored extramural (e.g. community-based, scientific society, etc.) leadership experiences. Opportunities in years three and four represent a partnership with the Mayo Clinic Graduate School of Biomedical Sciences (MCGSBS) Career Development Internship (CDI) program, whereby career exploration will be coupled with intentional leadership skills building/practice. A summary of planned activities is depicted in Figure 1. Year one activities are described in more detail below.

**Figure 1:**
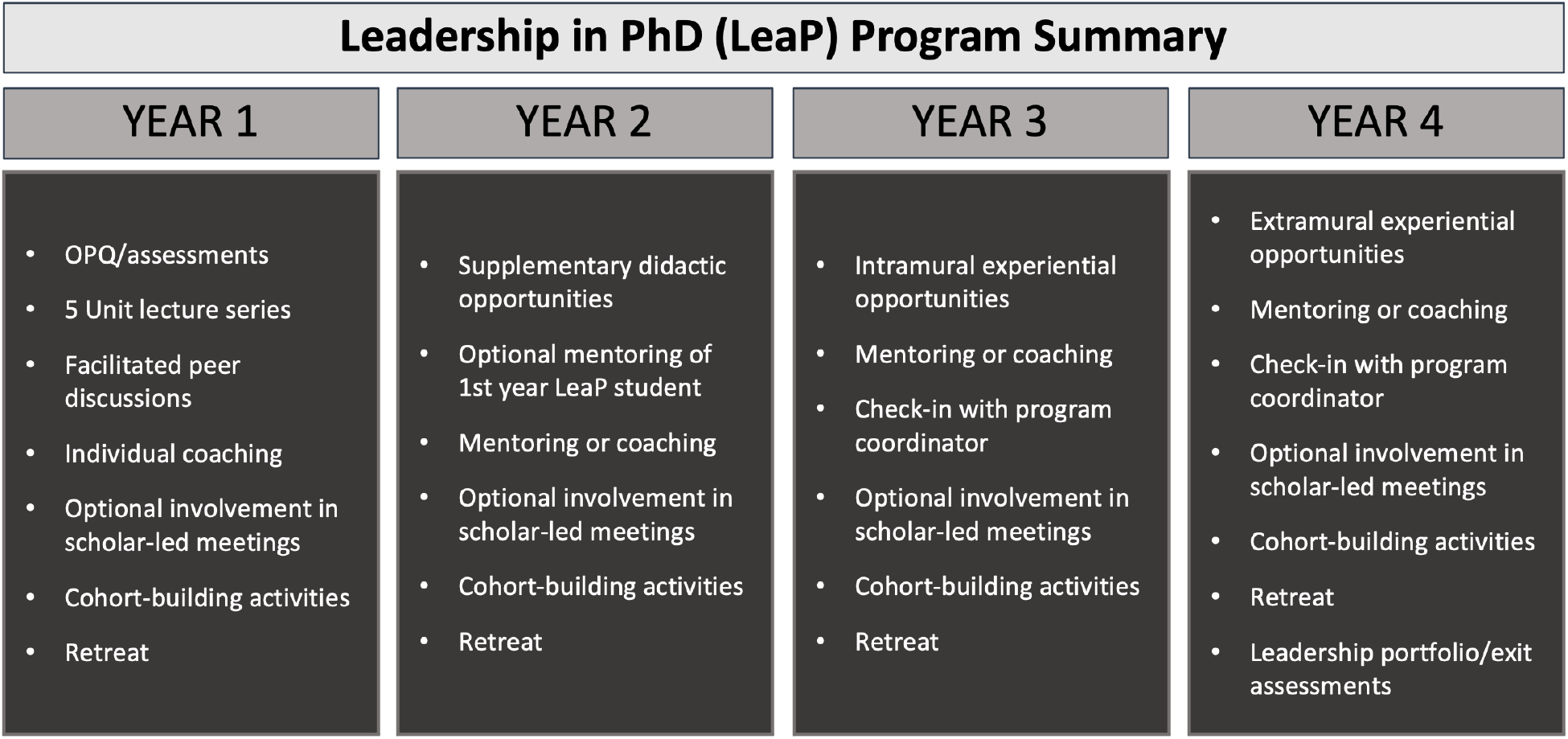
Program summary. Shown (L to R) are summaries of key activities proposed for each year in the program.

The first year was presented over nine months during the 2021-2022 academic year, with a staggered start with PhD coursework to give time for students to acclimate to their home campus, classes, and laboratory rotations. Prior to official program launch, an introductory session was held where program mission, motivation, and structure were shared and discussed. Five units were outlined each with sub-themes that were intentionally designed to address the ten leadership competencies described above (see competency map in Figure 2).Units were structured as monthly modules each containing a seminar (open to all graduate students), a facilitated discussion, and an individual coaching session. Two units (emotional intelligence and communication) were presented as two-month modules with six touchpoints instead of three. Most meetings were held virtually given that LeaP included MCGSBS students from all three Mayo Clinic campuses (Rochester MN, Scottsdale AZ, Jacksonville FL). Brief unit descriptions are as follows:

**Figure 2:**
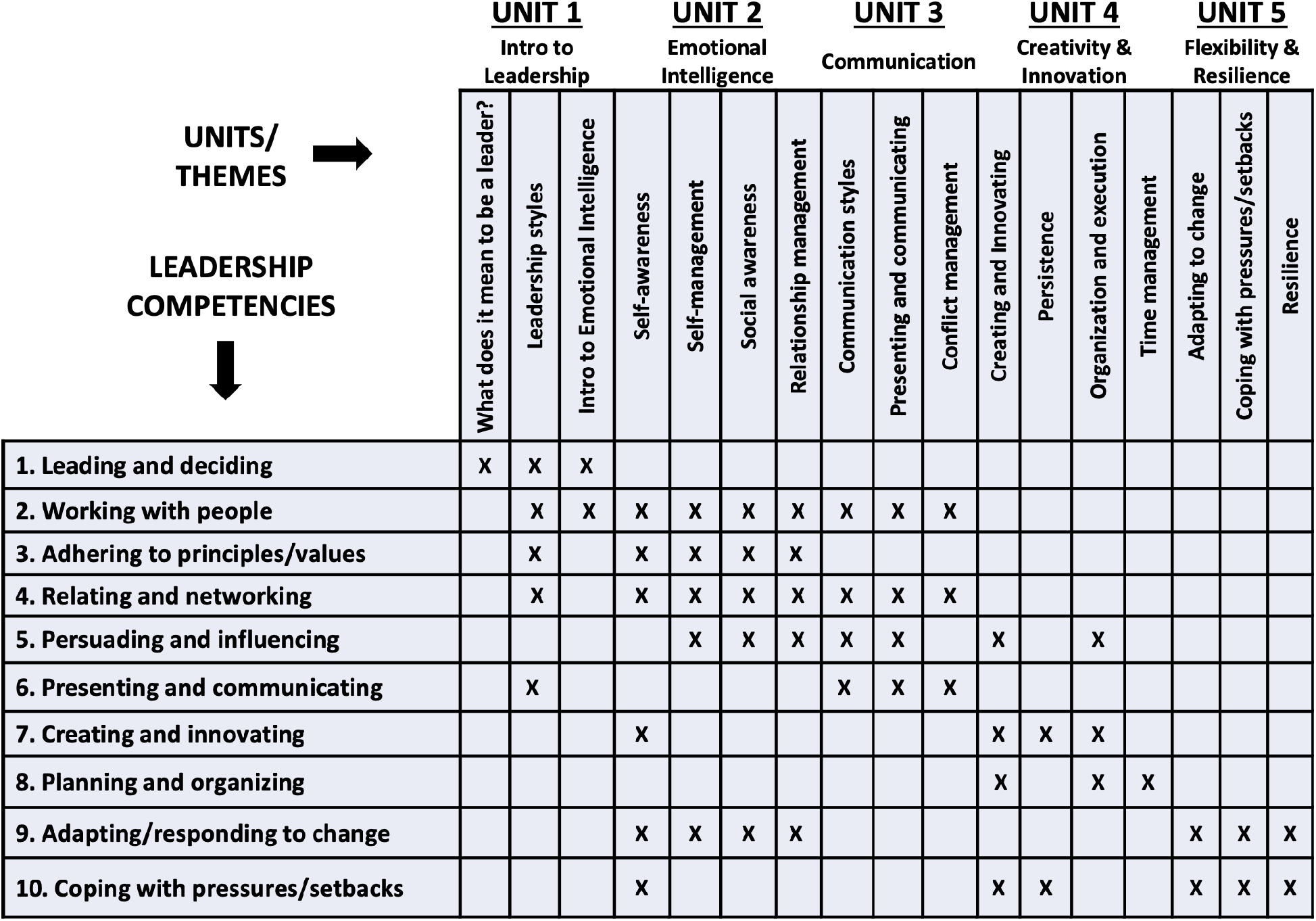
Competency map. Shown (across) are the five year one units and associated sub-themes. Rows indicate specific leadership competencies derived from the OPQ. An “X” indicates competency coverage addressed by a given unit sub-theme.

### Unit One: Introduction to leadership

Sub-themes: 1) meaning of leadership, 2) leadership styles, and 3) introduction to emotional intelligence. Discussion highlights: Students were asked to a) identify their go-to leadership styles, b) consider when/why it may be beneficial to adapt their leadership style, and c) reflect on how to go about navigating leadership styles while maintaining an inclusive and equitable environment.

### Unit Two: Emotional intelligence (EI)

Sub-themes: 1) self-awareness, 2) self-management, 3) social awareness, and 4) relationship management. Discussion highlights: a) the importance of (and strategies to develop) self-awareness social awareness, b) ways to practice self-management relationship management, and c) scenarios to prompt discussion around how to apply EI skills in the workplace.

### Unit Three: Communication

Sub-themes: 1) presenting and communicating, 2) communication styles, and 3) conflict management. Discussion highlights: a) communicating for impact, b) the importance of relationships, c) communication styles and how they relate to EI, and d) prompts to discuss real-life applicability (e.g. how does understanding of our own and others’ communication styles help us manage conflict?).

### Unit Four: Creativity and Innovation

Sub-themes: 1) creating innovating, 2) planning time management, 3) organization execution, and 4) persistence. Discussion highlights: a) brainstorming techniques, b) the importance and impact of creativity and innovation, and c) conversation around how to create a safe and supportive environment that encourages different points of view, experimentation, failure, and persistence.

### Unit Five: Flexibility and Resilience

Sub-themes: 1) adapting to change, 2) flexibility, 3) dealing with pressures and setbacks, and 4) resilience. Discussion highlights: flexibility and resilience strategies were shared during an integrated in-person two-day retreat held on the Rochester campus.

## 3 Evaluation Plan and Results

### Evaluation plan

LeaP program evaluation was developed collaboratively with the co-program directors and the OASES Director of Evaluation prior to the start of the program in Fall 2021. Being a new program, evaluation design focused on gathering both qualitative and quantitative data that would be useful for ensuring program quality and for identifying improvements for future years. Understanding participant views on the value of the sessions, as well as on the impact of the educational components and the program overall were top priorities.

After each of the five units, a survey was sent asking scholars to: a) rate the effectiveness of the session and coaching, b) describe 1-2 main take-aways from the discussions, c) rate the role of the lecture, discussion, and coaching in supporting those take-aways, and d) make suggestions for session and program improvements. The survey questions were uniform across all units but referenced unit-specific content. At the close of the program, qualitative data were gathered from participants in a focus group setting to query the value of the program and its various components, as well as opportunities for improvements.

Planned evaluation for years 2-4 includes post-session reaction surveys, periodic touchpoint surveys, year-end focus groups, peer assessments, PhD mid-program and exit surveys, and a final portfolio encompassing summaries and reflections on experiential (years 3 and 4) projects.A final OPQ will be administered and results discussed at individual exit meetings/coaching sessions. LeaP alumni will be encouraged to visit and periodically present to/serve on panels for current scholars. Following graduation, LeaP program directors will partner with MCGSBS alumni relations staff to maintain communication with and track career progression of former scholars.

### Results

Overall, the unit survey results were highly positive. Between 4 and 5 participants responded to each survey for a general response rate of between 50% and 63%. Participants were asked to rate their level of agreement (5-point Likert scale from strongly disagree to strongly agree) with various statements about the lecture, discussion, and coaching, as well as whether they were likely to use the information they learned in the session. Sessions were highly rated with all respondents across all topics either strongly agreeing or somewhat agreeing that: a) the lectures broadened their understanding of various leadership styles, b) the post-lecture discussions were effective in deepening their knowledge about the topic, c) coaching supported their personal understanding of their strengths and areas for growth, and d) they were likely to use the information they learned in the unit. In those ratings, most participants strongly agreed with those statements (Figure 3A). Notably, for all sessions, all respondents strongly agreed that the coaching was supporting their personal understanding of their strengths and areas for growth.

**Figure 3:**
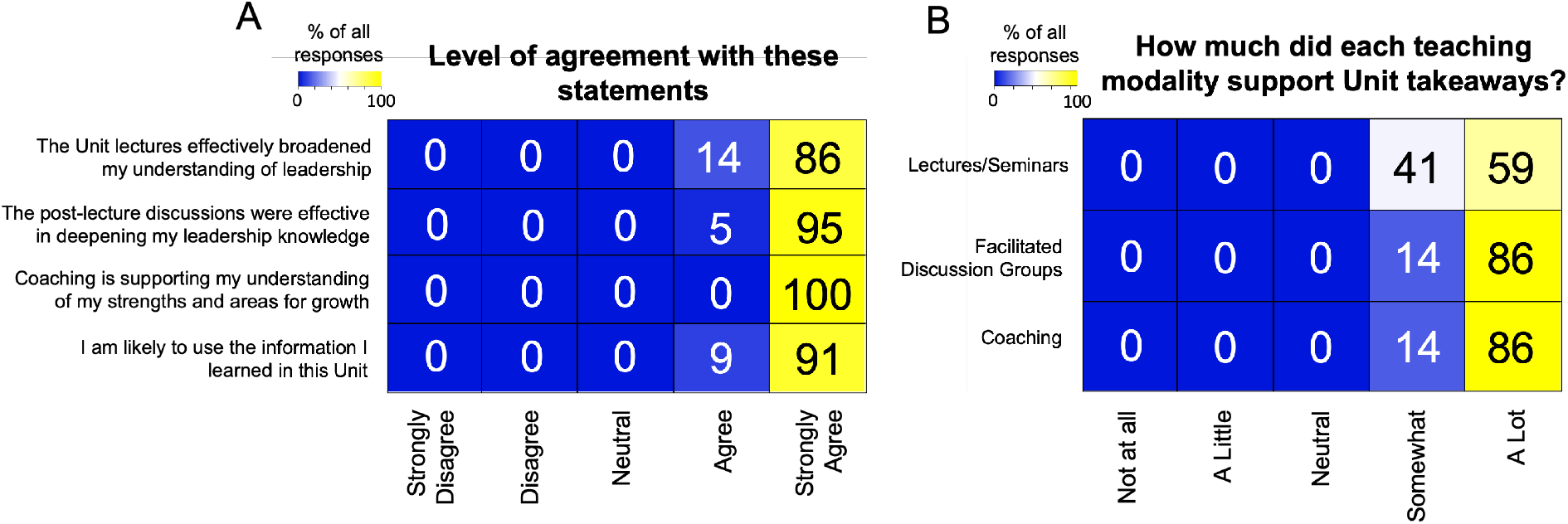
Aggregate unit survey data. (A) A heatmap depicting participant level of agreement with survey statements. (B) A heatmap depicting participant opinions regarding the three main teaching modalities. Data for both (A) and (B) are shown as a percentage of total responses across all five units.

For each unit, participants were asked about their main take-aways, as well as the extent to which the lecture, the discussion, and the coaching supported those take-aways. Participants all indicated that the lecture, discussion, and coaching supported their take-aways either “somewhat” or “a lot” with most checking “a lot” (Figure 3B). Participants had relatively few suggestions for improving the teaching modalities; however, suggestions that did surface tended to be related to creating more opportunities for participation and discussion.

In order to gather qualitative data on the value and impact of the program, a focus group was held in early June 2022 following completion of Year 1 of the program. Six of the eight participants attended the focus group. The two participants who were unable to attend the focus group provided feedback separately, one in an individual interview and the other with written responses. The same set of guiding evaluation questions was used for all participants:

1. Strengths: What were the strengths of the program in your opinion?
2. Group Dynamics: Describe the group dynamics of the participants in the LeaP program. Was the size of the group good (or too small or too large)? How important was it that all the participants come from groups underrepresented in biomedical research?
3. Facilitation: Tell me about how the co-program directors facilitated the group. To what extent did the way they facilitated work for you? Is there anything that could have been done to improve the facilitation?
4. Coaching: Tell me about the coaching. Was the coaching valuable to you? If yes, in what ways? Is there anything that could have been done to improve the coaching?
5. Impact: What impact has participation in the program had on you? Do you feel that you are able to apply what you learned in the program? What impact do you think participation in the program will have on you as you go through the rest of your PhD program?
6. Changes and Recommendations: Were the session topics the right ones? Are there any other topics that could or should have been included? What suggestions do you have for improving the program for the participants who start this upcoming academic year?
7. Other Comments: What other suggestions or comments do you have about the program that have not been covered in the questions above.

Participants were overwhelmingly positive about the LeaP program, its value, and its impact. Summarized, representative findings included:

- The size of the cohort was just right. A significantly larger cohort would have made it more difficult for participants to share openly and freely. That openness and ability to be vulnerable with the group helped participants reflect on and make sense of their experiences through the year without judgement and led to a greater sense of belonging.
- Participants in the program developed long-lasting bonds with each other over the year and with the program leads. Having come to the program with shared lived experiences, they were able to gain validation for their experiences as first-year PhD students.
- The coaching was clearly a critical component of the program. Participants described tailored experiences to their individual needs and styles. Participants appreciated session flexibility and the lack of a rigid agenda or approach. All described how they had changed over the year and been impacted by the experience.
- Participants all thought that the topics chosen were important, and the format of the units—with a presentation by a speaker on Tuesdays, a follow-up discussion among the participants, along with the coaching—helped them reflect on and think deeper about the topic and how it applied to their graduate school experience. The separation between the presentation and the follow-up discussion was appreciated, as it left time for participants to “marinate” on the topic.
- Participants noted repeatedly that the program co-directors did an excellent job facilitating the discussions, being prepared with probing questions, helping moderate when discussions got intense, and avoiding judgement. They also attributed the sense of belonging that they gained, in part, to their facilitation.
- Some suggested that it would be better if the individuals who present on Tuesday do not attend the discussion session on Thursday because their presence sometimes changed and restricted open discussion among the participants. Although the speakers only attended the discussion sessions once during the year, the participants noticed the difference in the quality of the discussion.
- Some longer-term impacts described by the participants included: a) having a cohort they could rely on throughout their PhD program, b) finding a sense of belonging, c) feeling validated through cohort discussions, d) building their communication skills so they could advocate for themselves and others, including about physical and mental well-being, e) gaining clarity about their personal and professional goals, f) learning to be assertive and intentional about those goals, and g) coming to see themselves as leaders and feeling empowered to lead.
- A few suggestions were made for improvements to the program: a) ensuring that the discussions held on the Thursday following the presentation are specifically for open discussion among the participants (without the presence of the speakers), b) getting the cohort together as early as possible in the academic year so that connections could be nurtured sooner, c) having the program be in person, d) developing a cohort mission statement within each group, e) ensuring a better gender balance among the cohort, f) expanding the program with additional cohorts, and g) keeping quality coaching as a central aspect of the program for future cohorts.
- Participants would like the program to continue for their cohort, with Year 2 involving the development of a project proposal that could impact their communities and with Year 3 being used to implement that project. They noted that the format of Year 1—with a focus on reflection and gaining great knowledge of their strengths and areas for growth—was an important stepping stone enabling them to have a broader impact in future years.

## DISCUSSION

Persistent concerns with representational diversity, equity and fairness, and feelings of inclusion/belonging in biomedical research underscore the need to diversify research leadership and support leadership skills development (23–26). LeaP tackles these issues by providing a space for underrepresented biomedical research trainees to build community, self-reflect, learn new leadership skills, and to progressively practice and refine those skills in a safe and supportive environment. To that end, LeaP joins a growing number of programs aiming to modernize biomedical education and training by providing early-stage leadership development opportunities (26–31). What sets LeaP apart from many of these programs, however, is that LeaP 1) promotes skills development early on in biomedical training, 2) intentionally weaves diversity, equity and inclusion topics into the curriculum, 3) cultivates social interactions via a cohort model that supports trainee persistence and retention (32), and 4) promotes not only the acquisition of new skills, but skill perdurance via coaching and longitudinal (four year) engagement. The value of LeaP derives from the intentional combination of these component parts.

Evaluations and feedback from participants, faculty, staff, and collaborators were overwhelmingly positive. That said, LeaP is not without limitations. Based on promising early returns, it is tempting to propose rapid program growth/scaling to make the program accessible to anyone who wants to participate. While this is an admirable goal, it will be important to carefully reflect on what made the program meaningful to participants in the first place. Two themes stood out in the evaluations: trust/the cohort effect and the value of coaching. Indeed, most participants cited a strong culture of trust – between each other as well as with the program directors – as critically important. So how could/can growth occur while maintaining a safe and supportive environment? Parallel cohorts is one idea, although issues of cohort variability would have to be addressed. This includes careful selection and onboarding of any new facilitators and coaches. A team/task-based approach may help alleviate this concern, whereby curricular components (facilitation, coaching, etc.) could be split into teams guided by a lead facilitator or coach that could help set and maintain standards. Scaling LeaP effectively will take time, thoughtful consideration, and input from/consultation with education and leadership experts.

It is important to note that we do not yet know the longterm impact of LeaP. How effective are such trainee-directed leadership programs? This question is difficult to answer given often inadequate long-term follow-up and widely varying definitions of what it means to be “successful” (24, 33). That said, with LeaP, we are anticipating lasting impact by leveraging a two-pronged approach consisting of 1) coaching to enhance self-reflection and accountability, and 2) experiential/service-learning opportunities to reinforce and amplify self-efficacy and personal growth (34, 35). Both tactics are evidence-based leadership development strategies that we believe could be implemented as a component of formal PhD/advanced degree training, particularly for historically marginalized and underrepresented trainees.

What opportunities lie ahead for LeaP? Certainly, there is potential to adapt the model to other health professional programs at Mayo Clinic. To that end, there are numerous leadership development initiatives at other medical institutions (7, 28, 30, 35–37) that could integrate core LeaP concepts/approaches to tailor the experience to medical students, clinical fellows, or other professional degree seekers. Additionally, while trainee-focused initiatives are highly valuable and critically important, efforts to “train the trainer” to best support UR trainees (38) are needed and could be an interesting expansion of/addition to LeaP offerings. Another version of this approach is the concept of reverse mentoring (39, 40). This experience could be particularly beneficial for LeaP participants as it would show them that learning and leadership are bi-directional and that while established faculty/leaders may have expertise in many areas, many are willing to seek out and engage in continued personal growth.

Finally, should a program like LeaP be available to both minority and majority trainees? While thinking about the answer to this question, it is important to remember and consider historical disparities in biomedical leadership as well as obstacles and challenges that underrepresented trainees continue to face today. Equity and fairness do not equate with equality and some disparities may need to be addressed with specialized programs such as LeaP, where a culture of trust and safety is paramount and may be compromised in open settings where “outsider” feelings persist. We opted to open some activities – notably the didactic seminars – to all graduate students, while the majority were targeted to LeaP scholars. Striking a balance of targeted and untargeted activities will require continuous feedback and reflection from all stakeholders moving forward. Ultimately, our hope is that the initial success of LeaP is recognized as a viable model of UR trainee leadership development, such that similar approaches are initiated, adapted, and sustained at biomedical institutions worldwide.

## ACKNOWLEDGEMENTS

The authors wish to thank the Kern National Network for Caring and Character in Medicine (KNN)/Mayo Clinic for generous financial support of this pilot initiative. We also thank Mayo Clinic Graduate School of Biomedical Sciences leaders, faculty, students and staff who participated in and served on planning/advisory committees. We further wish to acknowledge the content expertise provided by individuals within Mayo Clinic Workforce Learning and Leadership Development.

## REFERENCES

1. C. Woolston, Minority representation in US science workforce sees few gains. Nature 592, 805–806 (2021).

2. R. K. Fry, Brian; Funk, Cary (2021) STEM Jobs See Uneven Progress in Increasing Gender, Racial and Ethnic Diversity. (Pew Research Center).

3. D. Kozlowski, V. Larivière, C. R. Sugimoto, T. Monroe-White, Intersectional inequalities in science. Proceedings of the National Academy of Sciences 119 (2022).

4. W. M. Lambert et al., Career choices of underrepresented and female postdocs in the biomedical sciences. eLife 9 (2020).

5. J. L. Rosenbloom, M. Gumpertz, R. Durodoye, E. Griffith, A. Wilson, Retention and promotion of women and underrepresented minority faculty in science and engineering at four large land grant institutions. Plos One 12 (2017).

6. N. Chance, Exploring the Disparity of Minority Women in Senior Leadership Positions in Higher Education in the United States and Peru. Journal of Comparative International Higher Education 13, 206–225 (2021).

7. N. J. Brown, Promoting the success of women and minority physician-scientists in academic medicine: a dean’s perspective. Journal of Clinical Investigation 130, 6201–6203 (2020).

8. ACE (2017) American College President Study. (American Council on Education).

9. CUPA-HR (2020) 2020 Professionals in Higher Education Annual Report. (College and University Professional Association for Human Resources).

10. D. V. Y. Hunt, Lareina; Prince, Sara; Dixon-Fyle, Sundiatu (2018) Delivering through diversity. (McKinsey and Company).

11. M. R. Bond, A. E. Gammie, J. R. Lorsch, M. Welch, Developing a culture of safety in biomedical research training. Molecular Biology of the Cell 31, 2409–2414 (2020).

12. L. Canti, A. Chrzanowska, M. G. Doglio, L. Martina, T. Van Den Bossche, Research culture: science from bench to society. Biology Open 10 (2021).

13. A. M. K. Choi, J. E. Moon, A. Steinecke, J. E. Prescott, Developing a Culture of Mentorship to Strengthen Academic Medical Centers. Academic Medicine 94, 630–633 (2019).

14. S. J. Rockey, Mentorship matters for the biomedical workforce. Nature Medicine 20, 575–575 (2014).

15. T. Ahmed et al., MyNRMN: A national mentoring and networking platform to enhance connectivity and diversity in the biomedical sciences. FASEB BioAdvances 3, 497–509 (2021).

16. M. Ghee, D. Collins, V. Wilson, W. Pearson, The Leadership Alliance: Twenty Years of Developing a Diverse Research Workforce. Peabody Journal of Education 89, 347–367 (2014).

17. B. Tucker Edmonds et al., Diversifying Faculty Leadership in Academic Medicine. Academic Medicine Publish Ahead of Print (2022).

18. R. E. Boyatzis, Stimulating Self-Directed Learning Through the Managerial Assessment and Development Course. Journal of Management Education 18, 304–323 (2016).

19. D. Goleman, R. Boyatzis, A. McKee, Primal leadership. IEEE Engineering Management Review 37, 75–84 (2009).

20. S. Loeng, Self-Directed Learning: A Core Concept in Adult Education. Education Research International 2020, 1–12 (2020).

21. P. J. Smith, E. Sadler-Smith, I. Robertson, L. Wakefield, Leadership and learning: facilitating self-directed learning in enterprises. Journal of European Industrial Training 31, 324–335 (2007).

22. T. Joubert, N. Venter, “The Occupational Personality Questionnaire” in Psychological Assessment in South Africa. (2013), 10.18772/22013015782.25 chap. 20, pp. 277–291.

23. K. Elmassian, The growing void of leadership training in medical education. Mich Med 113, 28 (2014).

24. B. Onyura et al., Is postgraduate leadership education a match for the wicked problems of health systems leadership? A critical systematic review. Perspect Med Educ 8, 133–142 (2019).

25. M. W. True et al., Leadership Training in Graduate Medical Education: Time for a Requirement? Mil Med 185, e11–e16 (2020).

26. Y. C. Wang et al., Introducing the MAVEN Leadership Training Initiative to diversify the scientific workforce. Elife 10 (2021).

27. C. L. Byington et al., A Matrix Mentoring Model That Effectively Supports Clinical and Translational Scientists and Increases Inclusion in Biomedical Research: Lessons From the University of Utah. Acad Med 91, 497–502 (2016).

28. C. B. Meador et al., A workshop on leadership for senior MD-PhD students. Med Educ Online 21, 31534 (2016).

29. N. D. Spector, B. Overholser, Leadership and Professional Development: Sponsored; Catapulting Underrepresented Talent off the Cusp and into the Game. J Hosp Med 14, 415 (2019).

30. B. Kumar, M. L. Swee, M. Suneja, Leadership training programs in graduate medical education: a systematic review. BMC Med Educ 20, 175 (2020).

31. S. A. Blanchard, R. Rivers, W. Martinez, L. Agodoa, Building the Network of Minority Health Research Investigators: A Novel Program to Enhance Leadership and Success of Underrepresented Minorities in Biomedical Research. Ethn Dis 29, 119–122 (2019).

32. M. Estrada, Q. Zhi, E. Nwankwo, R. Gershon, The Influence of Social Supports on Graduate Student Persistence in Biomedical Fields. CBE Life Sci Educ 18, ar39 (2019).

33. E. T. Price et al., Are We Making an Impact? A Qualitative Program Assessment of the Resident Leadership, Well-being, and Resiliency Program for General Surgery Residents. J Surg Educ 77, 508–519 (2020).

34. K. R. Boehmer et al., Motivating Self-Efficacy in Diverse Biomedical Science Post-baccalaureate and Graduate Students Through Scientific Conference Implementation. Frontiers in Education 6 (2021).

35. J. A. Long et al., Developing leadership and advocacy skills in medical students through service learning. J Public Health Manag Pract 17, 369–372 (2011).

36. D. M. Blumenthal et al., Implementing a pilot leadership course for internal medicine residents: design considerations, participant impressions, and lessons learned. BMC Med Educ 14, 257 (2014).

37. C. Coe et al., Leadership Pathways in Academic Family Medicine: Focus on Underrepresented Minorities and Women. Fam Med 52, 104–111 (2020).

38. M. K. Norman et al., Delivering What We PROMISED: Outcomes of a Coaching and Leadership Fellowship for Mentors of Underrepresented Mentees. Int J Environ Res Public Health 18 (2021).

39. N. Garg, P. Singh, Reverse mentoring: a review of extant literature and recent trends. Development and Learning in Organizations: An International Journal 34, 5–8 (2019).

40. K. Gadomska-Lila, Effectiveness of reverse mentoring in creating intergenerational relationships. Journal of Organizational Change Management 33, 1313–1328 (2020).

